# Diffuse and regionally structured domestication of the common fig (*Ficus carica* L.) in the Mediterranean Basin

**DOI:** 10.1101/2025.09.18.677149

**Authors:** Bouchaib Khadari, Sanzhar Kakenov, Hafid Achtak, Jamal Charafi, Lamis Chalak, Sylvain Santoni, Finn Kjellberg, Amandine Cornille

**Affiliations:** AGAP Institut, University of Montpellier, CIRAD, INRAE, Institut Agro, Montpellier, France; Conservatoire Botanique National Mediterranéen, Antenne Occitanie - Languedoc-Roussillon, Montpellier, France; Division of Science, New York University Abu Dhabi, Saadiyat Island, Abu Dhabi, United Arab Emirates; University Cadi Ayyad, Polydisciplinary Faculty, Department of Biology, Safi Morocco; INRA, Regional Center of Meknes, Research Unit of Plant Breeding and Plant Genetic Resources Conservation, Meknès Morocco; Lebanese University, Faculty of Agronomy, Beirut, Lebanon; CEFE, CNRS, Univ Montpellier, EPHE, IRD, France

**Keywords:** *Ficus carica*, population genetics, domestication, gene flow, Mediterranean Basin, conservation genetics, perennial crops, breeding, local landraces

## Abstract

Understanding domestication in perennial crops is crucial for unraveling the evolutionary trajectories that have shaped crop genetic diversity and for guiding future conservation and breeding efforts. The domestication of perennial Mediterranean fruit trees is less studied than that of annual crops. The common fig (*Ficus carica* L.) is thought to have been domesticated in the Levant before dispersion across the Mediterranean. However, pre-human fossils of *F. carica* in Europe suggest an ancient wild presence, challenging the assumption of a single eastern origin. We used microsatellite markers to genotype 949 fig accessions comprising cultivated and spontaneous individuals collected from 14 Mediterranean and Near Eastern countries representing *F. carica sensu stricto* and its two close relatives, *F. carica* subsp. *rupestris,* and, *F. colchica*. Principal component analysis revealed that *F. carica sensu stricto* forms a cohesive genetic group distinct from its two relatives, which are actually different species. Bayesian clustering revealed three major gene pools within *F. carica sensu stricto* — Moroccan-Algerian, Northern Mediterranean, and Levantine—each containing both cultivated and spontaneous individuals. The Levantine group was the most differentiated, while the other two were closer genetically, together reflecting a longitudinal Mediterranean structure. Cultivated and spontaneous figs were genetically indistinguishable within each region, supporting a model of diffuse, regionally independent domestication rather than a single Levantine origin. These results highlight spontaneous populations and local landraces as key reservoirs of genetic variation, particularly in Morocco, Algeria, and the Levant. Our study provides a foundation for future genomic work to identify the genetic basis of key traits, emphasizing that practical breeding and conservation strategies should rely on regional biodiversity rather than a narrow set of elite cultivars to enhance fig resilience and productivity in the face of climate change and emerging disease pressures.

## Introduction

Crop domestication reflects the evolving codependence between plants and human societies (Purugganan, 2019). Understanding domestication strategies revolves around several key questions: the identification and geographical origin of wild progenitors, the genetic exchanges driving domestication, and their consequences on population genetic diversity and structure of the crop and its wild relatives, as well as the ultimate pace of domestication (Allaby et al., 2008; Fuller et al., 2023; Gaut et al., 2015; Meyer & Purugganan, 2013). Knowledge of genetic diversity, admixture, and population structure can also guide breeding programs aimed at conserving genetic units, which are crucial for ensuring food security (Purugganan & Jackson, 2021).

Fruit trees are vital crops in our food systems and offer models for understanding the anthropogenic pressures that have shaped the domestication of perennial plant species. Yet the domestication trajectories of fruit trees are considerably less investigated than those of annual crops such as maize (*Zea mays* L.), wheat (*Triticum aestivum* L.), and rice (*Oryza sativa* L.) (Cornille et al., 2014; Gaut et al., 2015; Meyer et al., 2012; Miller & Gross, 2011). Fruit tree species often have very long domestication histories, due to their long generation times, generally high levels of genetic variation, and high levels of gene flow among populations (Besnard et al., 2017; Cornille et al., 2014, 2019; Gaut et al., 2015; Meyer et al., 2012; Miller & Gross, 2011). These protracted trajectories may reflect centuries of cultivation, recurrent hybridization with local wild relatives, and continual re-selection for agronomically important traits (Besnard et al., 2018; Diez et al., 2015). In the Mediterranean region, the domestication history of trees is even less well documented than that of temperate fruit trees. Among them, the olive (*Olea europaea* L.), the grape (*Vitis vinifera*), and, more recently, the almond (*Prunus dulcis*) trees are the best studied. In contrast, others, such as fig and carob, have received far less attention. Genetic studies have revealed extensive gene flow and suggested the possibility of multiple domestication origins, yet, determining whether domestication occurred once or multiple times remains particularly difficult to resolve, largely because of the pervasive gene flow between wild and cultivated populations (Barazani et al., 2023; Besnard et al., 2017, 2018; Besnard & Rubio de Casas, 2016; Decroocq et al., 2025; Diez et al., 2015; Dong et al., 2023; Julca et al., 2020; Lougmani et al., 2025; Myles et al., 2011).

Despite its cultural and economic value, the domestication history of the common fig (*Ficus carica* L.), a long-lived dioecious species of the Moraceae family, is poorly documented. This knowledge gap reflects several factors, including the unclear geographical distribution of wild *F. carica* and its relatives, unresolved taxonomy within the *F. carica* species complex, and a scarcity of population-level genetic studies. *Ficus carica sensu stricto* comprises both cultivated trees, which are typically grown in orchards and clonally propagated, and spontaneous trees, derived from seedlings that arose by sexual reproduction between undomesticated individuals or between trees in orchards. Spontaneous populations occur in open or disturbed habitats, such as riverbanks, cliffs, and commensal niches around human settlements (e.g., walls, terraces with shallow water tables), and can become naturalized beyond their native range. In southern France, for instance, all cultivars are parthenocarpic, whereas this trait is rare in spontaneous populations along rivers. Fossil evidence from France and Italy (60,000–100,000 years before present) suggests a long-standing presence for figs north of the Mediterranean Sea, with leaf fossils from the Parisian region and stipitate fig fossils from Montpellier showing traits distinct from those of close *Ficus* relatives (Kjellberg et al., 2022; Saporta et al., 1876). The current distribution of spontaneous *F. carica sensu stricto* individuals is constrained mainly by the climate and by the range of its obligate pollinator, the fig wasp *Blastophaga psenes*, which must go through at least two generations per year to effect pollination and fruit production; the northern European limit of the fig wasp, near the 46th parallel in the 1980s, has recently extended northward into Germany due to global climate change (Rehberger et al., 2024). Additional spontaneous *F. carica sensu stricto* occurs mainly around the Mediterranean Basin and along the coasts of Libya, Egypt, and southern Israel, as well as in rare, favorable inland habitats. In eastern Mediterranean regions, such as Syria, it occurs west of the Jabâl al-Ansariya mountain range but not in the more arid interior (Kjellberg & Valdeyron, 1984).

Cultivated fig orchards are often located very close to spontaneous stands, creating opportunities for gene flow. Molecular data, particularly from simple sequence repeats (SSRs) and mitochondrial markers, indicate substantial local genetic continuity between cultivated and spontaneous populations across the Mediterranean and Middle East (Khadari et al., 2003, 2005; Oukabli et al., 2003), but the overall population genetic structure of *F. carica sensu stricto* (including both cultivated and spontaneous trees) is poorly resolved. In many parts of its range, *F. carica sensu stricto* also grows near closely related taxa that collectively form the so-called *Ficus palmata* (Forsk.) taxon, including *F. carica* subsp. *rupestris* (Boiss.), *F. colchica* (Grossh.), *F. johannis* (Boiss.), and *F. pseudosycomorus*. *Ficus palmata* extends from Ethiopia to India and Nepal, in climates from arid to subtropical humid, and is likely a species complex (Berg & Wiebes, 1992). In moist coastal regions along the Black Sea east of Trabzon, *F. colchica* grows on cliffs. By contrast, in drier areas of Anatolia, northern Syria, northern Iraq, and Iran, where no spontaneous *F. carica sensu stricto* individuals are found, cultivation of *F. carica sensu stricto* often occurs near natural populations of *F. carica* subsp. *rupestris*, the predominant form of fig trees in the Fertile Crescent (Khadari et al., 2003; Kjellberg et al., 2022). Spontaneous trees of *F. carica sensu stricto*, *F. carica* subsp. *rupestris*, and *F. colchica* may thus represent genuine wild lineages or introgressed populations, as gene flow between natural populations and cultivars is common in perennial fruit species (Aradhya et al., 2010; Besnard et al., 2018; Cornille et al., 2015; Delplancke et al., 2011; Flowers et al., 2019). Therefore, the relationships and distinctions between *F. carica sensu stricto*, *F. carica subsp. rupestris*, and *F. colchica* remain unclear, and the extent to which *F. carica subsp. rupestris*, and *F. colchica* to the *F. carica sensu stricto* gene pools is unknown. Archaeological interpretations have also led to confusion: for example, the male caprifig was once regarded as the wild ancestor of the female domesticated fig, although both sexes belong to the same species (Ikegami et al., 2024; Kjellberg et al., 1987). Claims of very early fig domestication from the Jordan Valley (Kislev et al., 2006, 2006) are unconfirmed, as they may represent examples of collected rather than cultivated fruits (Fuller & Stevens, 2019). Moreover, classical domestication indicators, such as larger seeds, do not apply to figs, as figs are selected for larger fruits and have more seeds rather than larger seeds. Therefore, the domestication history of the common fig remains to be fully reconstructed. As emphasized by (Zohary et al., 2012), evidence for early fruit tree domestication is largely circumstantial, and wild figs—like wild grapevine (*Vitis* sp.) and olives—have yet to be studied extensively at the population genetic level.

Here, we investigated the population genetic structure of figs using comprehensive SSR genotyping data obtained from cultivated and spontaneous trees sampled from 48 sites across 14 countries around the Mediterranean Basin, together with *F. carica* subsp*. rupestris* and *F. colchica*. Our study addresses a central question: Do cultivated fig genetic resources reflect a single origin of domestication in the Levant, followed by diffusion throughout the Mediterranean Basin, or do they reflect multiple, locally driven domestication events? To address this, question we asked five questions: (1) What are the major geographical axes of genetic structure among spontaneous and cultivated figs across the Mediterranean? (2) To what extent do locally cultivated and spontaneous figs resemble each other genetically? (3) Are there notable genetic hotspots that might contradict a diffuse, local domestication hypothesis? (4) Can we detect any genetic introgression from close relatives of *F. carica sensu stricto*? (5) Can we reject a model of domestication in the Levant, within the Fertile Crescent, followed by diffusion to other regions? By combining principal component analysis, individual-based Bayesian clustering, and spatial mapping of genetic diversity, we provide evidence for the local dynamics of domestication. We document three major gene pools within *Ficus carica sensu stricto* (Moroccan-Algerian, Northern Mediterranean Basin, and Levantine) and find no substantial genetic contribution from *F. carica* subsp*. rupestris* or *F. colchica* to *F. carica sensu stricto*.

## Results

### Fig trees form four main highly admixed populations, with spontaneous and cultivated plants grouped together

To explore the genetic structure of our fig samples, we first ran STRUCTURE on the complete dataset, which consisted of 884 individuals from *F. carica sensu stricto* (both spontaneous and cultivated), 27 *F. colchica* individuals, and 38 *F. carica* subsp*. rupestris* spontaneous trees. The *ΔK* statistic peaked at *K* = 4 (Supplementary Fig. S1). At *K* values above 4, we observed the same overall structure, with some potentially poorly defined additional clusters (Supplementary Fig. S1), likely representing fine substructure and noise rather than biologically significant groups defining different gene pools (Cullingham et al., 2020; Puechmaille, 2016).

To assess the robustness of the inferred structure, we repeated the analysis with the *F. carica* subsp*. rupestris* and *F. colchica* individuals excluded. Importantly, we obtained the same three clusters within *F. carica sensu stricto* (Supplementary Figs. S2b,c, S4, S5). We nevertheless visually examined the clustering pattern with higher *K* values to look for potential sub-structuring, but saw none. At large *K* values, *F. carica* subsp*. rupestris* appeared as a homogeneous, non-introgressed group, supporting its species-level distinction. Similarly, when *F. carica* subsp*. rupestris* was removed from the analysis with high *K* values, *F. colchica* emerged as a single, non-introgressed unit (Figs. S2c, S4). Based on these observations, we selected *K* = 4 as the most meaningful and interpretable level of structure in the gene pools of the dataset. Therefore, we used the full dataset results (884 *F. carica sensu stricto*, 27 *F. colchica*, and 38 *F. carica* subsp*. rupestris*) for further analyses.

STRUCTURE analysis revealed four major spatially resolved gene pools for *F. carica sensu stricto*, corresponding to distinct geographical regions: Morocco and Algeria (blue), referred to as MADZ hereafter; the Northern Mediterranean Basin (pink, MED); the Levant (orange, LVT); and South-East Anatolia (green, corresponding to *F. carica* subsp*. rupestris*, or RUP). The *F. colchica* samples from the southern coast of the Black Sea, from Trabzon eastward, were distinct but admixed with *F. carica and* subsp*. rupestris* and *F. carica sensu stricto*. Each cluster included both cultivated (white circles) and spontaneous (black circles) *F. carica sensu stricto* individuals (Fig. 1 and Supplementary Fig. S1). *F. colchica* individuals were assigned to specific gene pools but were highly admixed with the other gene pools based on their membership coefficients; indeed, only a few individuals were assigned to a single gene pool. Because the analysis lacked the power to identify this well-defined gene pool, we excluded *F. colchica* individuals from further analysis. By contrast, *F. carica* subsp*. rupestris* individuals showed strong assignment to a single cluster. We then examined the distribution of individual maximum membership coefficients (Supplementary Fig. S3). Most samples exceeded 0.90 for one cluster or another, justifying our choice of 0.90 as the full membership threshold. We considered 358 (37% of total) individuals as being admixed (i.e., with a membership coefficient < 0.90 in any cluster), most individuals showing genotypes intermediate between the MADZ and MED clusters (Table 1). Therefore, we identified four fig populations: three populations of *F. carica sensu stricto* (MADZ, MED, and LVTP), along with a single population of *F. carica subsp. rupestris* (RUP) (Fig. 1).

**Fig. 1.**
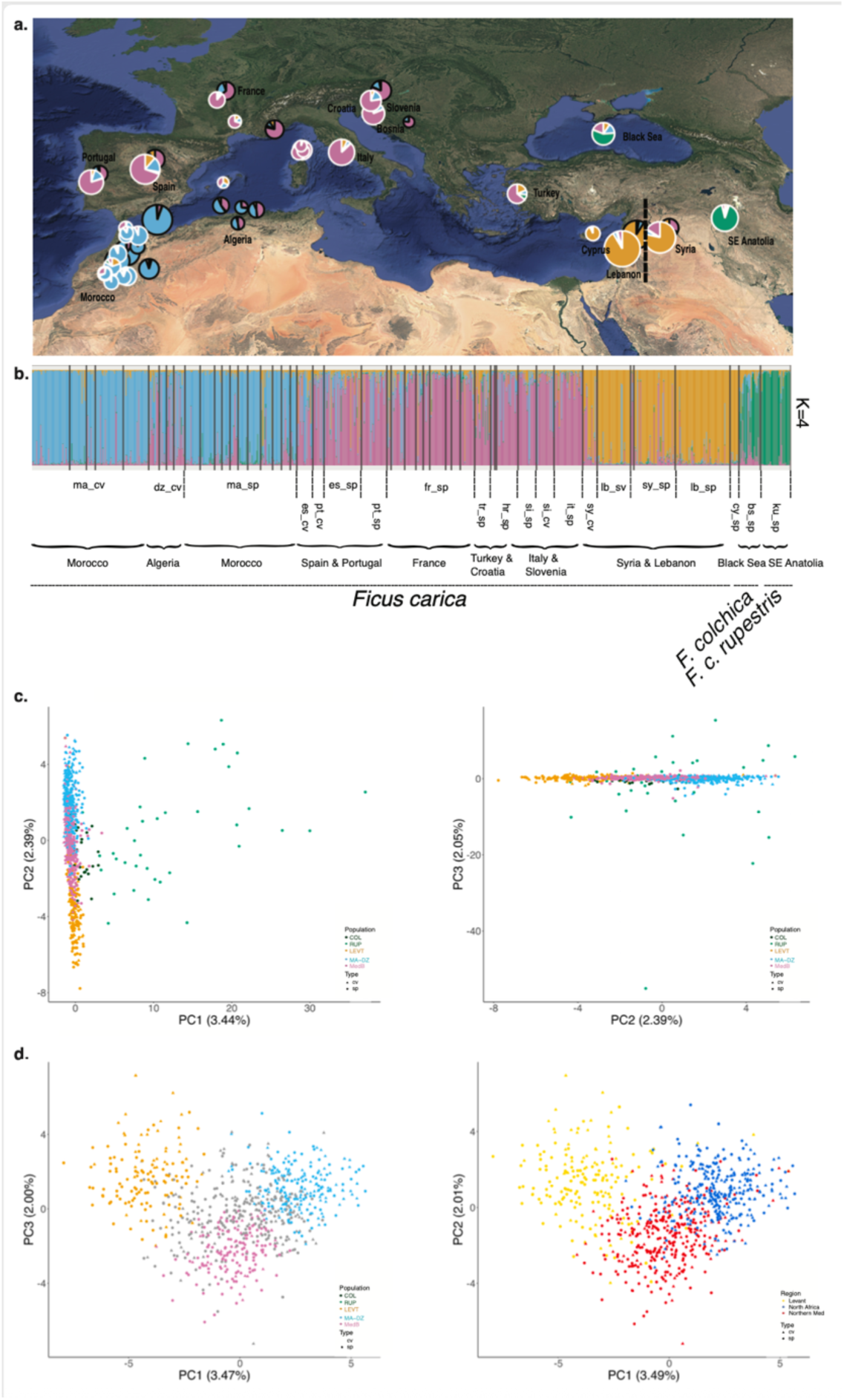
Population genetic structure and variation in cultivated and spontaneous fig trees from 14 countries, based on 14 SSR markers. This analysis is based on N = 949 individuals from *Ficus carica*, *F. carica* subsp. *rupestris*, and *F. colchica*. (a) Spatial genetic structure at *K* = 4, summarized as pie charts on the map. White-bordered pies denote spontaneous figs (sp), black-bordered pies denote cultivated figs (cv). Pie wedges show the relative average membership percentage in each genetic cluster. (b) STRUCTURE bar plot showing individual ancestry proportions at *K* = 4. Each vertical bar is a single fig tree; colors represent the inferred genetic clusters. (c) Principal component analysis (PCA) of the entire dataset of 949 individuals (*F. carica sensu stricto*, *F.* carica subsp. *rupestris,* and *F. colchica*). Two plots are shown: PC2 vs PC1 (left), and PC3 vs PC2 (right). Circles represent spontaneous (sp) individuals and triangles cultivated (cv). The PCA shows a clear separation of *F.* carica subsp. *rupestris* and *F. colchica* from *F. carica sensu stricto* along PC1, while *F. carica sensu stricto* displays a clear isolation-by-distance (IBD) pattern along PC2. (d) PCA restricted to *F. carica sensu stricto* only. Left, colors represent the genetic clusters inferred with STRUCTURE at *K* = 4; right, colors indicate the main geographical regions (Levant, North Africa, and Northern Mediterranean). Cultivated and spontaneous figs are genetically close within the three main clusters. Axis labels indicate the percentage of variance explained by each principal component.

In a principal component analysis (PCA), the first principal component (PC1) grouped almost all *F. carica sensu stricto* over a tiny range of values, while *F. carica* subsp*. rupestris* individuals were much more scattered along PC1. *F. carica* subsp*. rupestris* was genetically the most differentiated gene pool (Fig. 1), as validated by differentiation estimates (Table 2). When we removed *F. carica* subsp*. rupestris and F. colchica* individuals from the analysis, the three *F. carica* genetic clusters differentiated along an east-to-west continuum. This continuum was particularly marked for the Mediterranean and North Africa gene pools, while the Levant gene pool was more differentiated (Fig. 1c,d). Notably, spontaneous and cultivated figs within the Levant region formed one group in the PCA plot and exhibited very weak *F_ST_*, consistent with the distinct STRUCTURE gene pools. By contrast, cross-region comparisons (e.g., MADZ vs. RUP) were associated with the highest *F_ST_* values. We replotted the PCA results by coloring individuals according to their geographical origin instead of their gene pool. The results were largely congruent with those obtained when coloring based on PCA clustering (Fig. 1d).

These results confirm that *F. carica sensu stricto* constitutes a closely related genetic entity, separate from *F. carica* subsp*. rupestris*, and can be subdivided into three populations (MADZ, LVT, and MED). In addition, cultivated *F. carica sensu stricto* do not all cluster together, but instead cluster with their local spontaneous plants.

**Table 1.** Proportion of admixed individuals across fig genetic clusters inferred with STRUCTURE at *K*=4. Summary of the number (*N*ₐ_dₘ_) and proportion of admixed individuals (*P_adm_*) per species or subspecies *(F. carica sensu stricto*, *F. colchica*, and *F. carica* subsp. *rupestris*) in each STRUCTURE-inferred genetic cluster: Levant (orange, LVT), Morocco-Algeria (blue, MADZ), Northern Mediterranean Basin (pink, MED), and *rupestris* (green, RUP). *N*, total number of individuals per species or sub-species; *P*_ₐdₘ_, proportion of admixed individuals within each cluster (as percentage of the total admixed individuals for that taxon). Admixture was defined as assignment probabilities, 0.50<*Q-*values< 0.9, to any cluster.

| | Total $N$ | $N_{adm}$ | $P_{adm}$ | Orange Cluster<br>(LVT) | | Blue Cluster<br>(MADZ) | | Pink Cluster<br>(MED) | | Green Cluster<br>(RUP) | |
| --- | --- | --- | --- | --- | --- | --- | --- | --- | --- | --- | --- |
| | | | | $N_{adm}$ | $P_{adm}$ | $N_{adm}$ | $P_{adm}$ | $N_{adm}$ | $P_{adm}$ | $N_{adm}$ | $P_{adm}$ |
| *F. carica sensu stricto* | 884 | 323 | 36.5% | 52 | 16.1% | 104 | 32.0% | 163 | 50.5% | 4 | 1.2% |
| *F. colchica* | 27 | 24 | 88.9% | 1 | 4.2% | 4 | 17% | 2 | 8.3% | 17 | 70.8% |
| *F. carica* subsp.<br>*rupestris* | 38 | 11 | 28.95% | 0 | 0.0% | 0 | 0 | 1 | 9.1% | 5 | 45.45% |
| TOTAL | 949 | 358 | 37.7% |  |  |  |  |  |  |  |  |

**Table 2.** Genetic differentiation among the four fig populations inferred with STRUCTURE at *K* = 4. . The STRUCTURE analysis was based on *N=*596 individuals and 14 SSR markers. Populations were split into cultivated (cv) and spontaneous (sp) individuals to compare their genetic diversity. Colors correspond to the genetic groups identified by STRUCTURE at *K* = 4. Admixed individuals were excluded from this analysis. The values in the upper triangle represent Jost’s D values, while those in the lower triangle represent *F_ST_* values, indicating genetic differentiation between gene pools. All *P-*values are significant (*P* < 0.001).

|  | MADZ-cv | MADZ-sp | MedB-cv | MedB-sp | LEVT-cv | LEVT-sp | RUP-COL-sp |
| --- | --- | --- | --- | --- | --- | --- | --- |
| MADZ-cv | – | 0.01 | 0.12 | 0.12 | 0.23 | 0.25 | 0.31 |
| MADZ-sp | 0.01 | – | 0.13 | 0.12 | 0.24 | 0.26 | 0.28 |
| MedB-cv | 0.09 | 0.09 | – | 0.02 | 0.19 | 0.16 | 0.27 |
| MedB-sp | 0.09 | 0.08 | 0.01 | – | 0.18 | 0.14 | 0.26 |
| LEVT-cv | 0.18 | 0.17 | 0.13 | 0.13 | – | 0.04 | 0.33 |
| LEVT-sp | 0.18 | 0.17 | 0.10 | 0.09 | 0.03 | – | 0.30 |
| RUP-COL-sp | 0.16 | 0.13 | 0.11 | 0.12 | 0.15 | 0.14 | – |

**Fig. 2.**
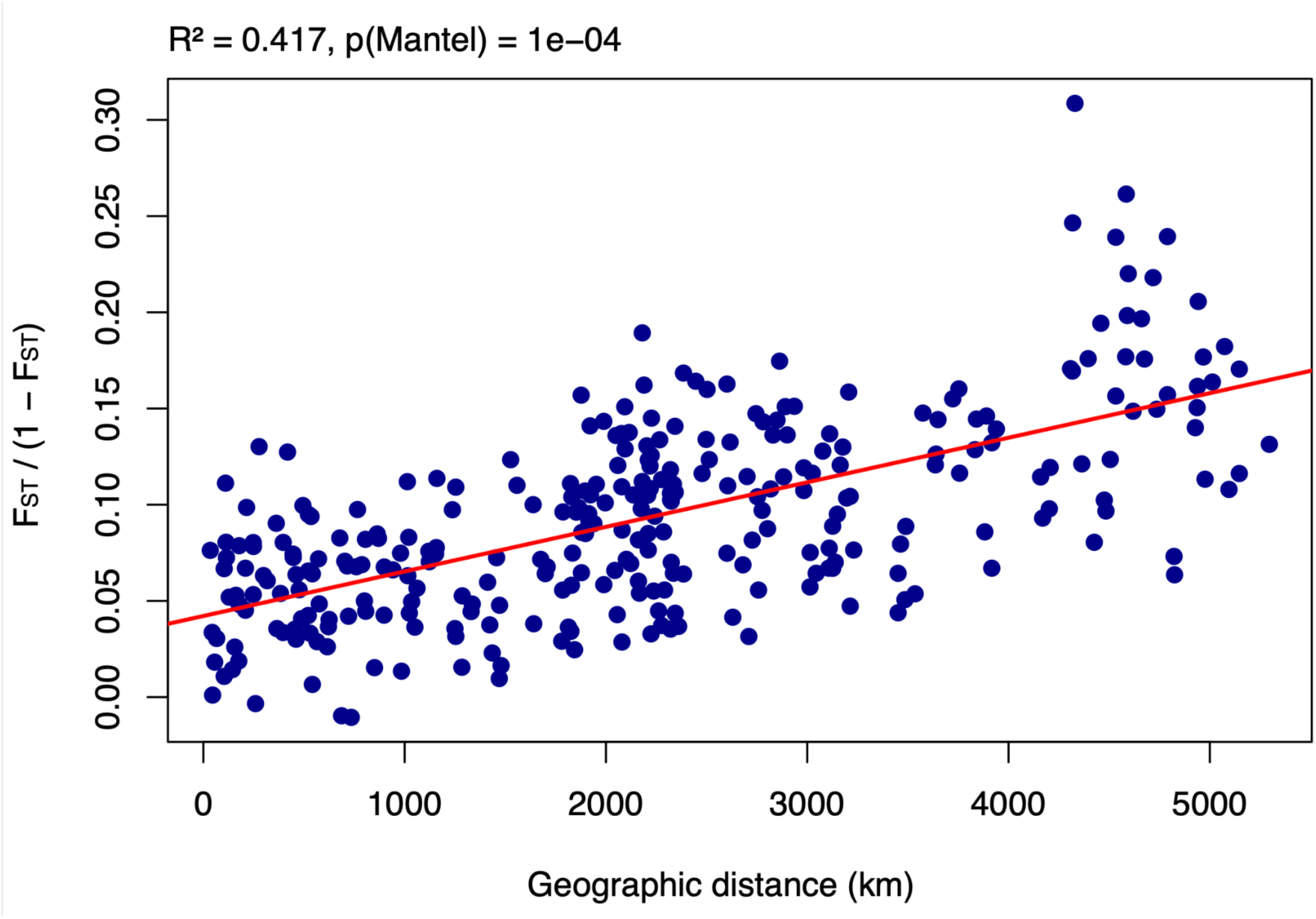
Isolation by distance (*F_ST_/*(*1−F_ST_*) as a function of geographical distance in spontaneous *Ficus carica sensu stricto* (*N* = 26 sites). Each blue circle represents a comparison between two geographical sites. The red line indicates the linear regression fit (intercept = 0.042, R^2^ = 0.417). Statistical significance for the Mantel test: *p* = 1 × 10^−4^.

### Spatial regional diversity and differentiation patterns suggest distinct regional histories

The *F. carica* subsp*. rupestris* population (RUP) showed the highest overall allelic richness (*A_R_* = 10.25) and private allelic richness (*Aₚ* = 4.58; Tables 3, S3, and S4). By contrast, private allelic richness was lower in the Northern Mediterranean Basin (MED) than in any other population. The MADZ and LVT populations exhibited medium to high diversity and retained more private alleles than the MED and RUP populations, as expected for populations at the range margins of a genetic continuum. Notably, the MADZ population showed moderate allelic richness (*A_R_* = 4.2–4.7 in cultivated forms). The LVT and MED populations also maintained substantial genetic diversity, with cultivated populations frequently exhibiting high observed heterozygosity (Table 3). In terms of inbreeding (Table 3), the *F_is_* values were overall low, generally below 0.10 or even negative, except for *F. carica* subsp*. rupestris* (0.125). Furthermore, all clusters maintained relatively high levels of allelic richness (Table 3), as well as observed heterozygosity, even in the cultivated groups.

We detected significant isolation-by-distance (IBD) within spontaneous *F. carica sensu stricto* as a whole (Fig. 2). Spatial genetic structure (SGS) within each spontaneous population was weak but significant (0.006 < *Sp* < 0.011), suggesting substantial historical gene flow within each population (Table 3). However, the LVT population showed the highest *Sp* values, suggesting that there were more historical barriers to gene flow in this population. This finding suggests that spontaneous tree populations are structured by geography, indicating barriers to gene flow among gene pools, despite having historically experienced massive gene flow within each population. An Estimated Effective Migration Surfaces (EEMS) analysis further revealed strong regional contrasts in effective migration rates, consistent with barriers to gene flow and regional divergence histories. Posterior probabilities of deviations from the average migration rate (log[m]) highlighted significantly smaller effective migration corridors (brown regions, P(log[m] < 0.05)) in the Levant and South-East Anatolia, suggesting limited recent gene flow with surrounding populations. Conversely, we detected higher migration corridors (blue, P(log(m) > 0.9)) between parts of the Western Mediterranean Basin and North African regions (Supplementary Fig. S7a). These patterns align with our STRUCTURE analysis, indicating limited connectivity between *F. carica sensu stricto* in the Levant and the two other gene pools. The log-posterior trace plot from the EEMS MCMC suggested that the chain has mixed well without strong trends (Supplementary Fig. S7b), indicating convergence of the algorithm and robustness of the inferred migration surface. In parallel, the posterior mean diversity rates (log[q]) reflected pronounced heterogeneity in local genetic diversity, with geographical diversity hotspots (positive log[q] values) identified in North Africa and parts of the Levant, consistent with the high private allelic richness and STRUCTURE-based genetic differentiation observed for these regions (Supplementary Fig. S7c). The posterior probabilities for deviations in diversity reinforced the significance of these spatial patterns (Supplementary Fig. S7d). Last, comparisons between observed and fitted genetic dissimilarities showed a global fit between model predictions and empirical data, although some variance remained unexplained, particularly over larger geographical distances (Supplementary Fig. S8). These results support the occurrence of IBD, leading to regional structuring of spontaneous fig populations, which may be attributed to geographical barriers to gene flow.

**Table 3.**
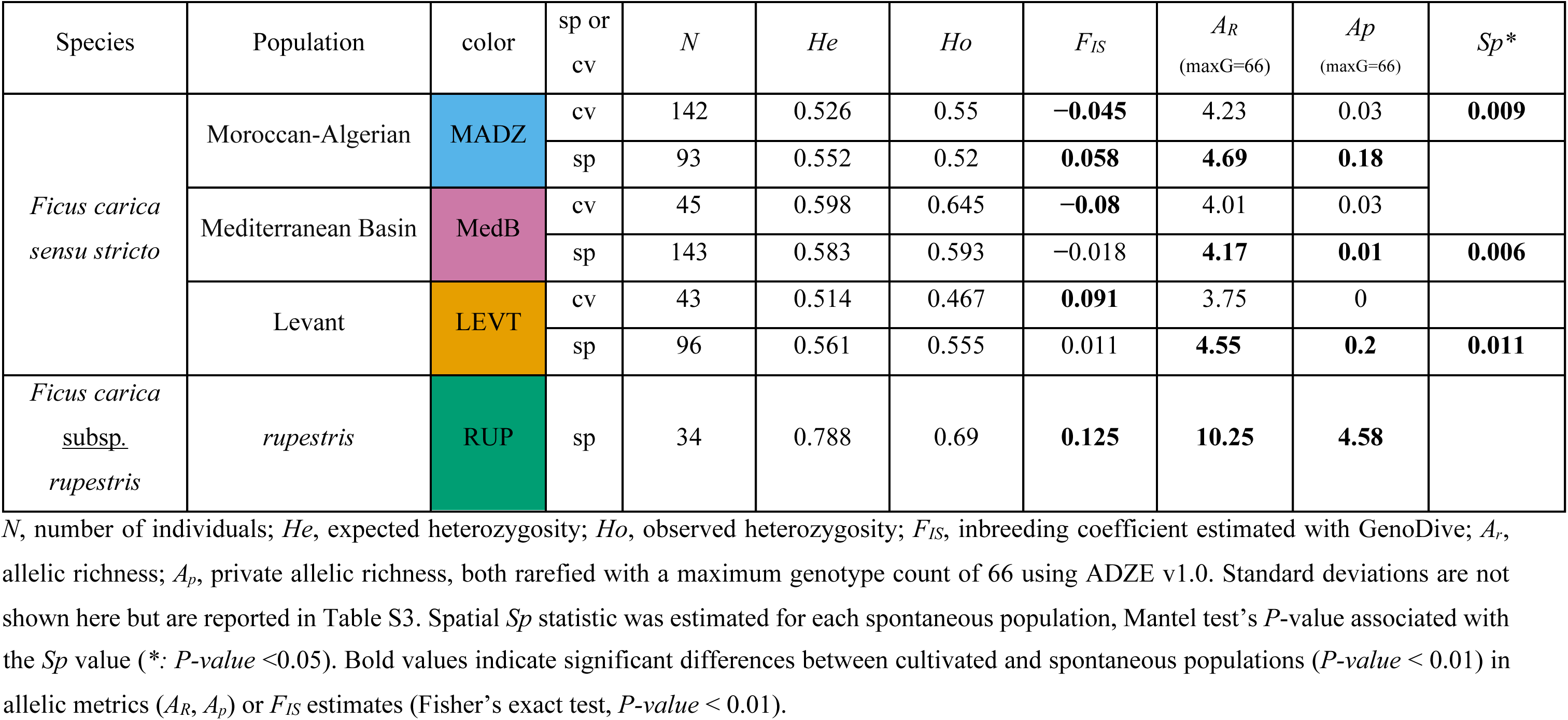
Genetic diversity estimates in four fig gene populations (cultivated or spontaneous) detected with STRUCTURE at *K* = 4. Admixed individuals, defined as those assigned to a genetic cluster with a membership coefficient <90%, were excluded from this analysis. Number of excluded individuals, *N*=98, including *F. colchica*; see Fig. 1 and Table 1. Cultivated (cv) and spontaneous (sp) individuals were analyzed separately within each population.

## Discussion

Our study provides the first broad-scale view of population genetic structure and diversity of cultivated and spontaneous *F. carica sensu stricto* across the Mediterranean Basin, together with a first glimpse into its relationship with *F. carica* subsp. *rupestris* y *F. colchica*. Using SSR markers, we identified three major genetic clusters from North Africa (Moroccan-Algerian), the Northern Mediterranean, and the Levant, each comprising both cultivated and spontaneous trees, and showing an east-to-west structured pattern. This pattern mirrors a common biogeographical feature in native perennial species of the region, reflecting long-term historical processes such as Pliocene–Pleistocene diversification and geographical barriers to gene flow. In most regions, cultivated and local spontaneous figs were genetically similar, suggesting frequent gene flow and/or local domestication from wild or naturalized populations. *Ficus carica* subsp. *rupestris* formed a distinct, well-defined cluster, whereas *F. colchica* appeared admixed with *F. carica sensu stricto*, indicating introgression from close relatives. However, denser sampling of the *F. palmata* complex (i.e., including *F. carica* subsp*. rupestris* and *F. colchica*) is needed to confirm this pattern. Together, these results challenge the hypothesis of a single domestication origin in the Levant followed by unidirectional diffusion. Instead, we propose a multi-regional domestication model in which local cultivated genotypes emerged repeatedly from regionally distinct spontaneous populations, with subsequent maintenance of genetic continuity between the two groups through pollen and seed flow. These findings have direct implications for fig breeding and conservation programs. Because cultivated and spontaneous trees within the same region share a large proportion of their genetic background, the most effective strategy for maintaining and enhancing fig genetic resources should be to prioritize local biodiversity, particularly that present in spontaneous and traditional landraces, over a narrow focus on a few elite cultivars. Breeding schemes that integrate locally adapted spontaneous trees can harness region-specific adaptive traits, such as drought tolerance, pest resistance, and fruit quality. Conservation programs should safeguard *in situ* populations as reservoirs of genetic diversity that underpin long-term resilience in Mediterranean fig cultivation.

### New insights into the *F. palmata* species complex: *F. carica* subsp*. rupestris* and *F. colchica* are distinct species

The PCA based on all individuals separated *F. carica* subsp. *rupestris* from *F. carica sensu stricto*; we observed the same separation for *F. colchica* and *F. carica sensu stricto*, although to a lesser extent. Almost all the diversity within *F. carica sensu stricto* was captured by the second principal component of the PCA. This pattern was confirmed by individual-based clustering analysis, which also distinguished *F. carica* subsp*. rupestris* and *F. colchica* from *F. carica sensu stricto*. We therefore conclude that *F. carica* subsp*. rupestris* and *F. colchica* differ from *F. carica* at the species level, and hereafter use the name *F. carica* only for *F. carica sensu stricto*. Likewise, *F. carica* subsp*. rupestris* is named *F. rupestris* hereafter. Two additional lines of evidence support the above conclusion: distinct natural habitats and the fossil record (Berg & Wiebes, 1992). Close relatives of *F. carica* are currently grouped under *F. palmata*, whose distribution extends from Ethiopia to India and Nepal, across climates ranging from arid to subtropical humid, suggesting that *F. palmata* is in fact a species complex (Berg & Wiebes, 1992). Within this group, *F. carica* subsp*. rupestris* is present in Anatolia, northern Syria, northern Iraq, and Iran, i.e., in drier areas where no wild *F. carica* populations are found (Khadari et al., 2003; Kjellberg et al., 2022). Each of these two species thus occupies a distinct natural habitat in separate geographical regions. Moreover, fossil evidence of *F. carica* in France and Italy, dating back 60,000–100,000 years, demonstrates a long-standing presence north of the Mediterranean. (Kjellberg et al., 2022; Saporta et al., 1876). Based on morphology and habitat, *F. colchica* is probably more closely related to *F. carica* than to *F. carica* subsp*. rupestris*, although definitive relationships will require the analysis of full genomic data. *F. colchica* grows under very moist conditions, whereas spontaneous *F. rupestris* individuals are found in dry regions outside the natural habitat of *F. carica*, although *F. rupestris* may be cultivated there.

### Three gene pools and regional domestication patterns in *F. carica*

Our genetic analyses revealed three main gene pools within the native range of *F. carica*: the Moroccan-Algerian (MADZ), Northern Mediterranean Basin (MED), and Levant (LVT) gene pools, following a common east-to-west Mediterranean phylogeographical structure observed in many native perennial species of the region. MADZ and MED were the closest gene pools, representing a broad western group that encompasses North Africa and the northern Mediterranean coast, from Portugal to Aegean Turkey. Cultivars from Algeria (spontaneous individuals were not collected) were somewhat intermediate, showing mixed assignment to MADZ and MED, which may reflect historical gene flow and local domestication, possibly linked to the colonial-era relationship between France and Algeria. By contrast, the LVT gene pool, representing the eastern Mediterranean, was clearly distinct from the two closely related western gene pools. Spontaneous trees within each gene pool exhibited a spatially distinct genetic structure and overall IBD, with the strongest structuring observed in the Levant, likely due to geographical barriers that restrict gene flow. This east-to-west genetic differentiation fits the pattern documented for many native Mediterranean tree species, which originated before the onset of the summer-dry climate in the Pliocene (∼3.2 million years ago [Mya]) and diversified into distinct eastern and western lineages (Feliner, 2014; Petit et al., 2005; Suc, 1984; Suc et al., 2018).

Within each gene pool, spontaneous and cultivated trees were genetically indistinguishable, indicating that domestication occurred locally rather than through large-scale dissemination of early domesticated varieties from the Levant, as was the case with the olive tree (Besnard et al., 2018). Notably, the Levant population, which includes both cultivars and spontaneous trees, remains the most differentiated from the other two *F. carica* populations, without any marked loss of genetic diversity, in contrast to non-native Mediterranean fruit species such as apricot (*Prunus armeniaca*), whose wild relatives occur in China and show substantial loss of diversity in cultivated ranges ((Bourguiba et al., 2012) but see (Groppi et al., 2021)). This observation suggests that fig domestication likely occurred independently in at least two Mediterranean regions: the Levant in the east, and a western/central area. This domestication was characterized by close genetic relationships between seed-propagated spontaneous trees and clonally propagated cultivated trees. This pattern differs from the domestication pattern seen with other Mediterranean native fruit trees, such as olives, for which the propagation mode by cutting and grafting of cultivated populations strongly shaped the genetic differentiation between wild and cultivated populations at both regional and local scales (Ben-Dor et al., 2024; Besnard et al., 2013; El Bakkali et al., 2025; Flowers et al., 2019; Gurbuz-Veral et al., 2018; Zunino et al., 2024). The inability to genetically distinguish between spontaneous and cultivated figs at the regional level, even in areas such as the Levant, raises the question of whether the domestication of the fig tree followed a uniquely diffuse and local trajectory, not seen in other Mediterranean fruit tree species.

### Fig has a relationship with humans that is unique among Mediterranean domesticated fruit trees

How do the patterns observed for *F. carica* compare to those of other Mediterranean domesticated fruit trees? Four main domesticated fruit tree species are native to the Mediterranean Basin: fig, olive, grapevine, and carob (*Ceratonia siliqua*). All four species exhibit clear genetic evidence of presence in both the western and eastern Mediterranean before the advent of agriculture.

Olive and grapevine share a broadly similar history, with a primary domestication in western Asia, followed by the spread of cultivated forms across the Mediterranean. Cultivated grapevine is genetically closest to *Vitis sylvestris* from western and central Asia and the Caucasus, and more distantly related to *V. sylvestris* from Turkey and regions further west and north. Its dispersal into Europe involved introgression from local *V. sylvestris*, while it followed a distinct southern colonization route in North Africa. Similarly, wild olive (*Olea oleaster*) shows two major cytoplasmic lineages: a western pool (Morocco to Greece) and an eastern pool (Turkey and the Levant). Most olive cultivars (90%) carry the eastern cytotype, with the remaining minority carrying western cytotypes, especially in the western Mediterranean (Besnard et al., 2013). Nuclear data confirm that cultivated olives are generally closer to eastern *O. oleaster* accessions, with western cultivars being genetically distinct from local western *O. oleaster* (Gros-Balthazard et al., 2019). Domestication occurred in the eastern Mediterranean, followed by westward introduction, with incorporation of some local wild genetic material through hybridization and backcrossing. Unlike grapevine, which shows separate northern and southern dispersal routes, olive spread along a single east-to-west route. Carob presents a third pattern (Baumel et al., 2021; Viruel et al., 2019). Wild and cultivated carob are only weakly differentiated, suggesting that they underwent local domestication to a large extent. Cytoplasmic data reveal two main geographical lineages: one dominant in Morocco, Spain, Portugal, and Sardinia, but also sporadically present to the east; and another prevailing from southern France eastwards. As with the fig tree, spontaneous and cultivated individuals are genetically similar within each region, although the phylogeographical patterns differ.

The contrasting phylogeographies of these four species can be partly explained by their differing tolerances to frost and drought. Grapevine is the most frost-tolerant, followed by fig, olive, and carob, in descending order of tolerance; drought tolerance is lowest in fig, greater in grapevine and olive, and highest in carob. *V. sylvestris* survived glaciations along the northern Mediterranean rim and in western Asia (Dong et al., 2023). For fig, likely glacial refugia existed around much of the Mediterranean, producing present-day spontaneous populations with gradual genetic variation across space but more pronounced differentiation in the Levant. Olive survival during glaciations was restricted to the southern parts of the northern Mediterranean rim, islands, and the southern shore (Besnard et al., 2013), resulting in north– south similarities due to recolonization from the south. Carob appears to have been largely eliminated from the northern continental Mediterranean during climatic oscillations, persisting mainly on islands and in southern refugia at the eastern and western ends of the basin (Viruel et al., 2019). In all four species, the arid coasts of Egypt and Libya limited east–west contact along the southern Mediterranean rim.

Olive and grapevine exhibit the most pronounced domestication traits, such as large fruits in their cultivars. Cultivated grapevine is monoecious and self-compatible, unlike its dioecious wild ancestor. Olive is monoecious but self-incompatible (Saumitou-Laprade et al., 2017), requiring compatible cultivars for pollination. Fig and carob differ markedly: both are dioecious, with blurred boundaries between spontaneous and cultivated gene pools. In fig, pollination is carried out by species-specific wasps that develop in functionally male trees (traditionally called *F. carica caprificus*). By contrast, female trees (*F. carica domesticus*) produce sweet edible fruits, whether spontaneous or clonally propagated. Wild (spontaneous) female trees yield figs that are very similar to those of cultivars, and fruit production relies on either parthenocarpic varieties or systematic pollination by nearby male trees, thereby maintaining strong genetic continuity between wild and cultivated populations. Carob is propagated from seedlings, with male trees grafted with female scions to ensure pod production, while some males are preserved for pollination. As carob is more often grown for livestock fodder than for human consumption (Benmahioul & others, 2011), selection for fruit traits has been limited.

Among these four species, fig is unique in its commensal relationship with humans. The genetic continuity between local cultivars and spontaneous plants is consistent with observations from Morocco, where locally cultivated genotypes vary among villages, likely due to recruitment from nearby spontaneous trees (Achtak et al., 2010). In regions where *F. carica* grows in the wild but cultivation is limited, female trees derived from seeds germinating in field margins or ditches are often preserved and tended (Kjellberg et al., 2022). We propose a simple domestication pathway: wild figs are palatable and comparable in quality to cultivated forms; they naturally colonize rocky riverbanks but readily establish in human-made habitats such as villages and stone enclosures. Spontaneous trees producing desirable fruits are preserved, cared for, and occasionally propagated. Seed dispersal by humans consuming figs may have further encouraged their establishment in villages. We propose that *F. carica* became an early human commensal, with diffuse, long-term mass selection gradually fostering domestication traits in these ruderal populations. This ecological dynamic naturally produced genetic structures consistent with multilocal domestication.

### Implications for the conservation and sustainable use of local genetic resources

This study sheds new light on the genetic structure and domestication history of figs, with important implications for the conservation and sustainable use of local genetic resources. The exceptional diversity of *F. carica* subsp*. rupestris* and the unique alleles found in spontaneous populations from the Levant and Morocco-Algeria identify these wild (spontaneous) groups as valuable reservoirs of potentially adaptive genetic variation. Traits such as drought tolerance, disease resistance, and fruit quality may be preserved within these gene pools, underscoring their relevance for future breeding programs and climate-resilient agriculture.

Conservation strategies should prioritize both *in situ* preservation of genetically diverse spontaneous populations and *ex situ* collections. Particular attention should be given to regions where diversified fig orchards exist, such as those described by (Achtak et al., 2010), as these traditional orchards harbor high cultivar diversity and genetic continuity with spontaneous trees. Natural stands, or those that persist with minimal disturbance, may harbor unique genetic lineages shaped by local histories. In addition, in a global context where food products increasingly carry labels emphasizing their geographical origin, often through mechanisms such as Protected Designations of Origin (PDO), the concept of *terroir* has become central to both marketing and consumer perception. Our findings strengthen the biological and historical foundations of this trend by demonstrating that many local or regional fig landraces originated from, and remain embedded within, distinct regional genetic pools. This finding reinforces the cultural and agronomic significance of these varieties and provides a genetic basis for labeling policies that seek to valorize local landraces. While deeper genomic analyses will further refine our understanding, the present work already presents a strong argument for integrated policies that link genetic conservation, local identity, and product valorization, especially in light of the persistent gaps in *ex situ* representation identified by (Ramirez-Villegas et al., 2022).

Beyond applied perspectives, figs also provide an original model for advancing basic research on perennial domestication. Whole-genome studies can uncover the genomic basis of key domestication traits by identifying traces of selection, adaptive introgression, and demographic histories across Mediterranean gene pools. Such work will help clarify how domestication unfolded in long-lived, clonally propagated species and will identify candidate genes and genomic regions directly relevant for breeding resilient cultivars. By linking local biodiversity conservation with insights into the genomic architecture of domestication, figs offer a bridge between fundamental evolutionary research and the applied goal of developing climate-resilient, high-performing fruit trees.

## Materials and methods

### Sample collection and DNA extraction

Samples from a total of 949 fig (*Ficus* sp.) individuals were collected from 48 sites across 14 countries (see Table S1, Fig. 1 for detailed site descriptions, sample sizes, and geographical coordinates). The samples consisted of three morphologically distinct lineages: *F. carica* L*. sensu stricto* (*N* = 884), the wild fig *F. colchica* Grossh. (*N* = 27), and *F. carica* subsp*. rupestris* (*N* = 38). Based on morphology, *F. colchica* was considered the most divergent form within *F. carica,* while *F. carica* subsp*. rupestris* was considered to belong to the *F. palmata* species complex (Berg & Wiebes, 1992). Each site contained either the spontaneous type (wild type, labeled “Sp”), the locally cultivated type (“Cv”), or both fig types. The sampling sites spanned various geographical regions to capture genetic diversity across both spontaneous and cultivated populations (Table S1).

For *F. carica* L. (*N* = 884), samples were collected from 14 countries, including Lebanon (*N* = 111), Syria (*N* = 70), and Morocco (*N* = 198) (Table S1). Samples of *F. colchica* L. (*N* = 27) originated from the southern coast of the Black Sea, from Trabzon eastward. The third form, *F. carica* subsp*. rupestris* L. (*N* = 38) was sampled from sites in southeastern Anatolia (Diyarbakir, Elazig, Malatya). This comprehensive sampling ensured a robust representation of genetic diversity across both domesticated and spontaneous fig populations, facilitating an in-depth study of domestication history.

Plant tissue (leaves or shoots) was collected from each tree between 1985 and 1989, immediately dried, and stored at room temperature until DNA extraction. All samples were collected before the implementation of the Nagoya Protocol (October 2014), which ensures compliance through material transfer agreements (MTAs) when possible and applicable.

### DNA extraction, microsatellite amplification, and genotyping

Genomic DNA was extracted from 200 mg of dried leaves using a DNeasy Plant Mini Kit (QIAGEN, part # 69106) according to the manufacturer’s instructions, with the following modification: 1% (w/v) of polyvinylpyrrolidone (PVP 40,000) was added to buffer AP1. DNA quality and concentration were evaluated through spectrophotometry, spectrofluorometry, and agarose gel electrophoresis.

The following microsatellite markers were used for genotyping: 4B12, 6H2, F46B2, F4E9, TO6A12, TO6DO8, 6E2, and 4E12 (Achtak et al., 2009; Khadari et al., 2001), as well as the SSR markers LMFC24, LMFC30, LMFC28, LMFC32, LMFC26, and LMFC34 developed by Giraldo et al. (2005). All microsatellite loci were amplified using a PCR-based protocol adapted from Khadari et al. (2001). Each 20-μL reaction mixture contained 10 mM Tris-HCl (pH 8.3), 50 mM KCl, 2 mM MgCl₂, 10 pmol of each primer (with the forward primer labeled with a fluorescent dye), 200 μM of each dNTP, 1 U of Taq polymerase (KAPA Biosystems, part # KK1015), and 50 ng of template genomic DNA. The PCR conditions were as follows: an initial denaturation at 94 °C for 5 min; 30 cycles of 94 °C for 30 sec, 55 °C for 45 sec, and 72 °C for 1 min; followed by a final extension at 72 °C for 7 min.

Genotyping was performed using the automated capillary sequencer ABI PRISM 3130XL Genetic Analyzer (Thermo Fisher Scientific, Applied Biosystems), with GeneScan 500LIZ as the size standard (Applied Biosystems, part # 4322682). Alleles were scored using GENEMAPPER v.4.0 software (Applied Biosystems).

Only genotypes with less than 1% missing data per marker were retained. The frequency of null alleles was estimated using Genodive at each locus (Meirmans & Van Tienderen, 2004). All markers showed low rates of null alleles and were therefore all retained (Table S2).

### Population differentiation and genetic structure among wild and cultivated figs, and genetic diversity estimates

The genetic structure of the 949 fig genotypes was investigated by principal component analysis (PCA) to determine whether *F. carica sensu stricto*, *F. carica* subsp*. rupestris*, and *F. colchica* represented distinct genetic entities. The PCA was conducted using the dudi.pca function from the adegenet R package (Jombart & Ahmed, 2011).

Bayesian clustering, implemented in STRUCTURE v.2.3.3 (Pritchard et al., 2000), was then used to assign individuals to specific gene pools. This method estimates ancestry proportions from *K* hypothetical clusters while minimizing deviations from the Hardy-Weinberg equilibrium and linkage disequilibrium. *K* values ranging from 1 to 15 were tested, with 15 independent runs for each *K* value. Each run involved 50,000 burn-in steps followed by 500,000 Markov chain Monte Carlo (MCMC) iterations.

STRUCTURE analyses were conducted using either the entire dataset (*N*=949) or excluding *F. carica* subsp*. rupestris* samples and then excluding both *F. carica* subsp*. rupestris* and *F. colchica* samples. These additional analyses helped us estimate how removing these *F. carica* subsp*. rupestris* and *F. colchica* affected the clustering patterns within typical *F. carica*. The same parameters were used as described in the previous paragraph.

To determine the most likely number of clusters (*K*), the *ΔK* method (Evanno et al., 2005) was used, as implemented in STRUCTURE HARVESTER (Earl & vonHoldt, 2012). Since *ΔK* identifies the strongest but not necessarily the finest population structure (Puechmaille, 2016), the STRUCTURE bar plots were visually examined for other values of *K*. After selecting the best *K* value, an individual was assigned to a particular cluster if its assignment probability to that cluster was greater than 90% (see results). This threshold was chosen based on the distribution of membership coefficients inferred with STRUCTURE for each cluster (see results below), to ensure that each identified cluster contained well-assigned individuals and was not solely made up of potentially admixed genotypes. To summarize and visualize the STRUCTURE outputs, the R package pophelper v.2.3.0 (Francis, 2016) was used. A population was then defined as a group of individuals with a membership coefficient below 0.90 to a given cluster by using the rationale of a panmictic group. Individuals with coefficients below 0.90 were considered “admixed.” This comprehensive approach enabled the identification of the major genetic groups and finer-scale population structure within our dataset.

Three distinct methods were used to further explore genetic variation and differentiation among the STRUCTURE-detected genetic groups. First, a PCA was performed, restricting the data to *F. carica sensu stricto*. In the PCA plot, individuals assigned to a specific cluster with a membership coefficient of 0.90 or higher were color-coded according to their respective population. By contrast, admixed individuals (membership coefficient < 0.90) were shown in gray. Second, population relationships were examined by generating a neighbor-joining (NJ) tree (Huson, 1998; Huson & Scornavacca, 2012) using Nei’s standard genetic distance (Nei, 1987). This distance was calculated between pairs of individuals or populations (i.e., a group of individuals with membership coefficients ≥ 0.90 to a given cluster inferred with STRUCTURE) using Populations 1.2.31 (https://bioinformatics.org/populations/). The pairwise *F_ST_* between populations (Weir & Cockerham, 1984) was computed using GenoDive (Meirmans & Van Tienderen, 2004).

Descriptive genetic diversity values were calculated for each population identified by the STRUCTURE analysis. Allelic richness (*A_R_*) and private allelic richness (*A_P_*) were determined using ADZE (Szpiech et al., 2008) with a standardized sample size of *N_ADZE_* = 33 (Gmax = 66, representing the two chromosomes of one individual at each allele), which corresponds to the lowest observed count across populations. Observed and expected heterozygosity, as well as Weir and Cockerham F-statistics, were calculated and tested for deviations from the Hardy-Weinberg equilibrium in each population using GenoDive (Meirmans & Van Tienderen, 2004).

### Spatial pattern of diversity and differentiation

Diversity hotspots were investigated in cultivated and spontaneous figs by visualizing the spatial variation in allelic richness (*A_R_*) using QGIS (Quantum GIS, GRASS, SAGA GIS) with inverse distance weighting interpolation. This analysis was restricted to 46 collection sites with at least five successfully genotyped individuals, i.e., excluding sites 7 (yt_cv) and 11 (ba_sp). Three methods were used to further investigate spatial genetic structure, clines, and connectivity patterns. First, isolation-by-distance (IBD) was assessed by running a linear regression analysis between multilocus *F*_ST_/(1–*F*_ST_) estimates and distance for all pairs of sites using R and the ggplot2 package (Wickham, 2016). The slope, which expresses the degree of genetic structuring, was tested using 10,000 permutations on location, which is equivalent to performing a Mantel test. Pairwise *F_st_* values and geographical distances computed with GenoDive (Meirmans & Van Tienderen, 2004) were used to generate a scatterplot and calculate the Pearson’s correlation coefficient between genetic and geographical distance. Second, clines were investigated in the spontaneous populations by estimating the extent of spatial genetic structure (SGS) using the software SPAGeDI v1.3 (Hardy & Vekemans, 2002). SPAGeDI was applied to those individuals within each *F. carica sensu stricto* and the spontaneous populations, for which precise individual GPS estimates were available. The estimator of the kinship coefficient proposed by Nason (Loiselle et al., 1995) was used for pairs of individuals (*Fij*). *F_ij_* values were regressed on the spatial distance between individuals, *d_ij_*, and its natural logarithm, ln(*d_ij_*), providing the regression slopes *b_d_* and *b_Ld_*, respectively. Standard errors (SE) were calculated by jackknifing data over each locus. To assess whether SGS better matched predictions of IBD in two dimensions (*i.e.*, kinship decreasing approximately linearly with the logarithm of the distance), spatial positions of individuals were permuted 10,000 times (similar to a Mantel test) to test for SGS. The *Sp* statistic was then calculated, defined as *Sp*=– b(*Ld)*/(1−*F_N_*), where *F_N_* is the mean *F_ij_* between neighboring individuals, which was approximated by *F*(*d*) for the first distance interval (*dij*<1,000 m) (Hardy et al., 2006; Vekemans & Hardy, 2004). Connectivity patterns among fig populations were further investigated, applying the Estimated Effective Migration Surfaces (EEMS) method (Petkova et al., 2016). EEMS estimates how observed genetic dissimilarities deviate from an IBD model, producing maps of effective migration rates (m) and diversity (q) across a geographical landscape.

## Author contributions

AC, BK, FJ: conceptualization, methodology, investigation, data curation, formal analysis, visualization, writing – original draft, writing – review & editing, funding acquisition, project administration.

SK: methodology, investigation, data curation, writing.

SS, HA, JC, and LC data acquisition.

## Data availability

The microsatellite genotyping dataset in GenAlEx format is available on Zenodo: <u>xxxx/zenodo.xxxxxxx</u>. Scripts used for data analysis are deposited in the project’s GitHub repository: https://github.com/CornilleEclecticLab/Fig-SSR/.

