## Supplementary material for "Diffuse and regionally structured domestication of the common fig (*Ficus carica* L.) in the Mediterranean Basin"

**Table S1. Sample-level metadata and microsatellite allele sizes for 949 fig samples from 48 sites across 14 countries.** The table provides comprehensive metadata for each fig sample: sample type (cultivated or spontaneous), site and population codes, geographical origin, and microsatellite (SSR) allele sizes for 14 loci. The loci analyzed were 4B12, 4E12, 6H2, F46B2, F4E9, TO6A12, TO6D08, 6E2, LMFC24, LMFC30, LMFC28, LMFC32, LMFC26, and LMFC34. Each row represents an individual sample, detailing its genetic profile based on allele sizes at these loci.

See Excel file.

**Table S2.** Expected heterozygosity ($H_E$), observed heterozygosity ($H_O$), inbreeding coefficient ($F_{IS}$), and estimated null allele frequency (by Brookfield equation) for each genetic marker used in the study. The loci analyzed were 4B12, 4E12, 6H2, F46B2, F4E9, TO6A12, TO6D08, 6E2, LMFC24, LMFC30, LMFC28, LMFC32, LMFC26, and LMFC34.

| **Locus** | ***Ho*** | ***He*** | ***Fis*** | One-sided *p*-value | Proportion of null alleles (NA) |
|---|---|---|---|---|---|
| 4B12_A1 | 0.729 | 0.693 | −0.052 | 0.027 | 0.03 |
| 4E12_A1 | 0.655 | 0.664 | 0.013 | 0.28 | 0 |
| 6H2_A1 | 0.463 | 0.481 | 0.036 | 0.16 | 0 |
| F46B2_A1 | 0.5 | 0.489 | −0.023 | 0.211 | 0.01 |
| F4E9_A1 | 0.648 | 0.638 | −0.016 | 0.259 | 0.01 |
| TO6A12_A1 | 0.57 | 0.582 | 0.022 | 0.193 | 0 |
| TO6DO8_A1 | 0.539 | 0.49 | −0.1 | 0.006 | 0.03 |
| 6E2_A1 | 0.62 | 0.644 | 0.038 | 0.071 | 0 |
| LMFC24_A1 | 0.504 | 0.469 | −0.073 | 0.022 | 0.02 |
| LMFC30_A1 | 0.822 | 0.808 | −0.018 | 0.153 | 0.01 |
| LMFC28_A1 | 0.668 | 0.608 | −0.099 | 0.002 | 0.04 |
| LMFC32_A1 | 0.407 | 0.402 | −0.012 | 0.351 | 0.00 |
| LMFC26_A1 | 0.303 | 0.294 | −0.032 | 0.13 | 0.01 |
| LMFC34_A1 | 0.53 | 0.474 | −0.118 | 0.001 | 0.04 |
| **Average over loci** | 0.568 | 0.552 | −0.029 | 0.001 | 0.01 |

**Table S3.** Site-level summary of sample sizes and genetic diversity indices ($A_R/A_P$) for 48 fig collection sites. Each row represents a group (site or region) with associated counts and diversity metrics.

| **Site number** | **Site ID** | ***N*** | $H_O$ | $H_E$ | $F_{IS}$ | *P*-value | ***Ar* (Allelic richness)** | ***Ap* (Private allelic richness)** | $P_{(NA)}$ **(Proportion of null alleles)** |
|---|---|---|---|---|---|---|---|---|---|
| 1 | ku_sp | 38 | 0.69 | 0.78 | 0.12 | **0.00** | 5.13 | 1.44 | −0.18 |
| 2 | sy_cv | 18 | 0.47 | 0.47 | 0.00 | 0.49 | 2.73 | 0.01 | 0 |
| 3 | sy_sp | 52 | 0.57 | 0.57 | −0.01 | 0.37 | 3.22 | 0.03 | 0.01 |
| 4 | lb_cv | 42 | 0.49 | 0.51 | 0.05 | 0.06 | 2.83 | 0.09 | −0.02 |
| 5 | lb_sp | 69 | 0.55 | 0.56 | 0.02 | 0.16 | 3.16 | 0.05 | −0.01 |
| 6 | bs_sp | 27 | 0.58 | 0.60 | 0.02 | 0.28 | 3.26 | 0.11 | −0.01 |
| 8 | cy_sp | 11 | 0.58 | 0.51 | −0.15 | **0.00** | 2.89 | 0.07 | 0.08 |
| 9 | tr_iz_sp | 20 | 0.53 | 0.54 | 0.02 | 0.32 | 3.01 | 0.03 | −0.01 |
| 10 | ba_cv | 5 | 0.66 | 0.56 | −0.18 | **0.02** | 2.93 | 0.00 | 0.13 |
| 12 | hr_sp | 27 | 0.61 | 0.56 | −0.08 | **0.02** | 2.91 | 0.01 | 0.05 |
| 13 | si_sp | 23 | 0.58 | 0.54 | −0.07 | **0.04** | 2.90 | 0.00 | 0.05 |
| 14 | si_cv | 22 | 0.62 | 0.60 | −0.03 | 0.22 | 3.01 | 0.02 | 0.03 |
| 15 | it_sp | 36 | 0.60 | 0.56 | −0.07 | **0.01** | 2.96 | 0.00 | 0.05 |
| 16 | fr_ve_sp | 8 | 0.56 | 0.54 | −0.05 | 0.31 | 2.93 | 0.03 | 0.03 |
| 17 | fr_pn_sp | 20 | 0.52 | 0.54 | 0.03 | 0.29 | 2.86 | 0.01 | −0.02 |
| 18 | fr_fr_sp | 7 | 0.66 | 0.56 | −0.19 | **0.01** | 3.05 | 0.00 | 0.13 |
| 19 | fr_fo_sp | 11 | 0.56 | 0.56 | 0.00 | 0.44 | 2.98 | 0.00 | 0 |
| 20 | fr_pr_cv | 18 | 0.61 | 0.60 | −0.01 | 0.43 | 3.17 | 0.07 | 0.01 |
| 21 | dz_db_cv | 13 | 0.60 | 0.54 | −0.11 | **0.05** | 2.80 | 0.02 | 0.07 |

| | | | | | | | | | |
|---|---|---|---|---|---|---|---|---|---|
| 22 | dz_to_cv | 9 | 0.58 | 0.52 | −0.11 | 0.06 | 2.84 | 0.00 | 0.07 |
| 23 | dz_bo_cv | 9 | 0.57 | 0.50 | −0.14 | **0.02** | 2.88 | 0.01 | 0.07 |
| 24 | fr_so_sp | 9 | 0.48 | 0.57 | 0.17 | **0.01** | 3.01 | 0.00 | −0.1 |
| 25 | dz_cv | 14 | 0.63 | 0.55 | −0.15 | **0.01** | 2.99 | 0.00 | 0.1 |
| 26 | ba_i_sp | 5 | 0.56 | 0.53 | −0.05 | 0.30 | 2.64 | 0.00 | 0.03 |
| 27 | fr_br_cv | 17 | 0.55 | 0.58 | 0.05 | 0.15 | 3.06 | 0.00 | −0.03 |
| 28 | fr_sp | 14 | 0.58 | 0.55 | −0.04 | 0.25 | 2.86 | 0.03 | 0.03 |
| 29 | es_cv | 20 | 0.55 | 0.53 | −0.03 | 0.31 | 2.91 | 0.04 | 0.02 |
| 30 | es_sp | 47 | 0.55 | 0.57 | 0.02 | 0.22 | 3.03 | 0.02 | −0.01 |
| 31 | ma_rif_cv | 48 | 0.55 | 0.49 | −0.12 | **0.00** | 2.67 | 0.00 | 0.06 |
| 32 | ma_to_cv | 21 | 0.57 | 0.57 | 0.00 | 0.52 | 3.00 | 0.04 | 0 |
| 33 | ma_ta_sp | 19 | 0.51 | 0.50 | −0.02 | 0.40 | 2.70 | 0.00 | 0.01 |
| 34 | ma_nc_cv | 11 | 0.58 | 0.56 | −0.04 | 0.27 | 2.88 | 0.00 | 0.03 |
| 35 | ma_so_sp | 18 | 0.56 | 0.58 | 0.04 | 0.23 | 3.10 | 0.04 | −0.02 |
| 36 | ma_ch_sp | 9 | 0.60 | 0.55 | −0.10 | 0.07 | 2.98 | 0.00 | 0.06 |
| 37 | ma_mo_sp | 10 | 0.45 | 0.51 | 0.12 | **0.05** | 2.77 | 0.00 | −0.06 |
| 38 | ma_tu_sp | 10 | 0.58 | 0.54 | −0.08 | 0.14 | 2.87 | 0.01 | 0.05 |
| 39 | ma_ti_sp | 12 | 0.60 | 0.60 | 0.00 | 0.48 | 3.16 | 0.06 | 0 |
| 40 | ma_ou_sp | 16 | 0.55 | 0.58 | 0.05 | 0.13 | 3.04 | 0.02 | −0.03 |
| 41 | ma_cp_cv | 35 | 0.55 | 0.51 | −0.09 | **0.01** | 2.68 | 0.00 | 0.05 |
| 42 | ma_tb_sp | 15 | 0.52 | 0.56 | 0.07 | 0.13 | 3.02 | 0.04 | −0.04 |
| 43 | ma_se_sp | 11 | 0.53 | 0.52 | −0.02 | 0.40 | 2.85 | 0.00 | 0.01 |
| 44 | ma_nw_cv | 11 | 0.57 | 0.53 | −0.07 | **0.03** | 2.77 | 0.00 | 0.04 |
| 45 | ma_bh_sp | 8 | 0.54 | 0.54 | 0.00 | 0.51 | 2.84 | 0.06 | 0 |
| 46 | ma_hl_sp | 32 | 0.53 | 0.51 | −0.04 | 0.31 | 2.84 | 0.01 | 0.02 |

| | | | | | | | | | |
|---|---|---|---|---|---|---|---|---|---|
| 47 | pt_cv | 14 | 0.58 | 0.57 | −0.01 | 0.42 | 2.94 | 0.00 | 0.01 |
| 48 | pt_sp | 32 | 0.67 | 0.58 | −0.16 | **0.00** | 2.86 | 0.03 | 0.12 |

In Bold: values for which P-value are significant (P<0.05)

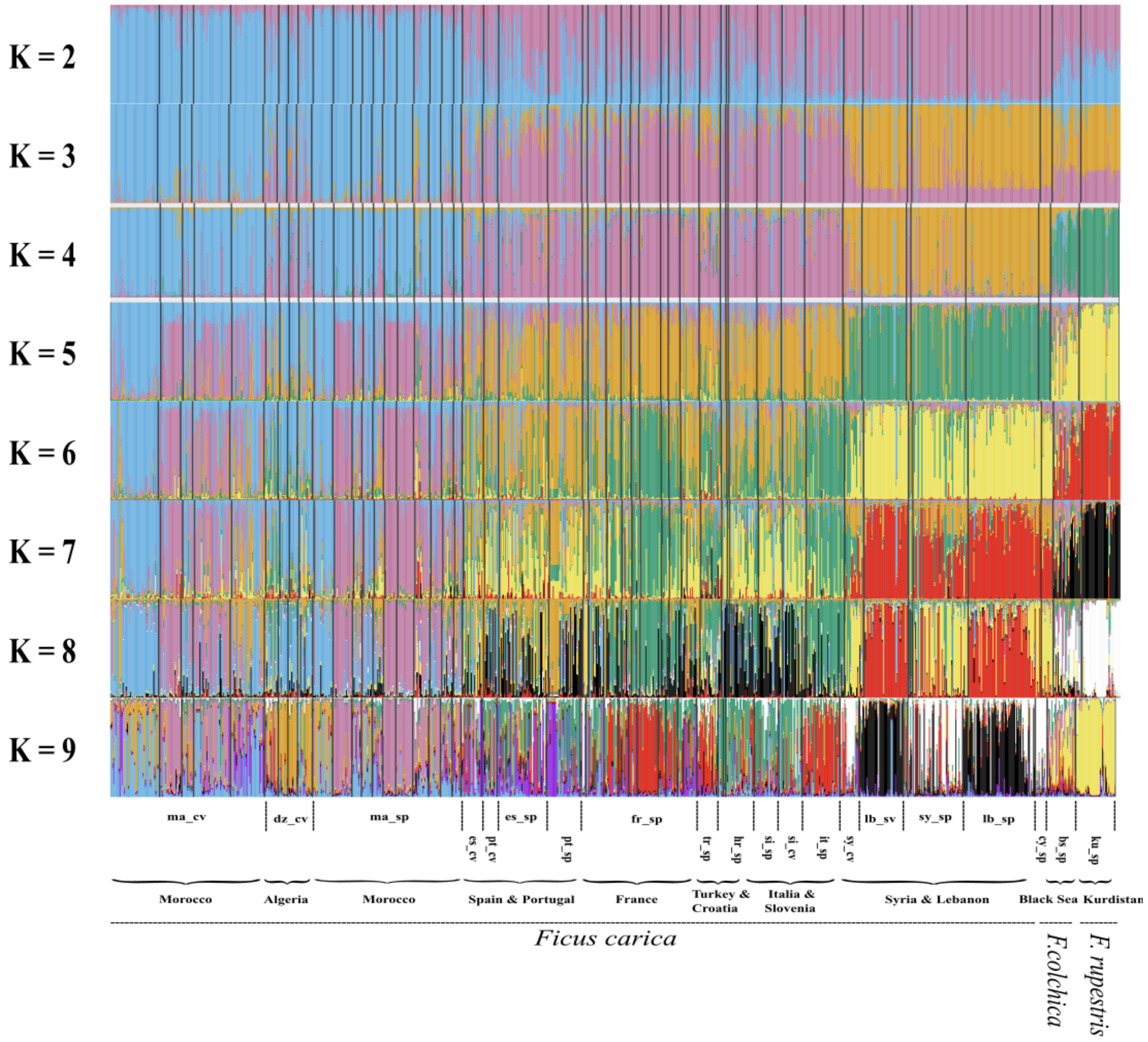


**Figure S1. Population structure and admixture among spontaneous and cultivated fig trees inferred from *K* = 2 to K = 9 with STRUCTURE.**

Each individual is represented by a single vertical bar partitioned into K-colored segments, and each vertical bar corresponds to the proportion of an individual's genome belonging to a given cluster. When multiple clustering solutions (modes) arose among replicate runs, the proportion of simulations supporting each mode is indicated. This analysis was carried out on 949 *Ficus* genotypes comprising 884 *F. carica sensu stricto*, 38 *F. carica* subsp *rupestris*, and 27 *F. colchica* individuals.

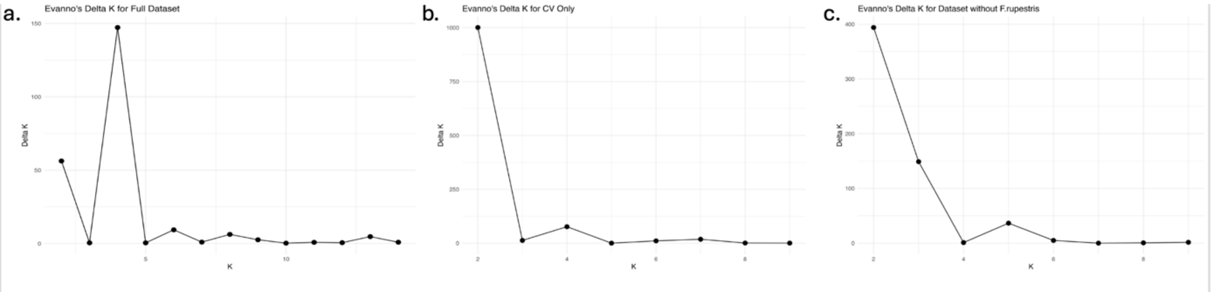


**Figure S2. Evanno's ΔK statistic for determining the optimal number of genetic clusters (K) in different datasets for the cultivated and spontaneous fig trees used to choose the most likely K value for STRUCTURE analysis.**

The ΔK statistic was estimated with Structure Harvester. STRUCTURE results are based on 14 microsatellite markers. (a) Full dataset consisting of all *Ficus* samples (N=949). (b) Dataset excluding *F. carica subsp rupestris*. (c) Dataset comprising only cultivated individuals, excluding *F. carica subsp. rupestris* and spontaneous (wild) individuals (i.e., bk). Each plot shows the *ΔK* values for different K values, with peaks indicating the most supported number of genetic clusters for each dataset.

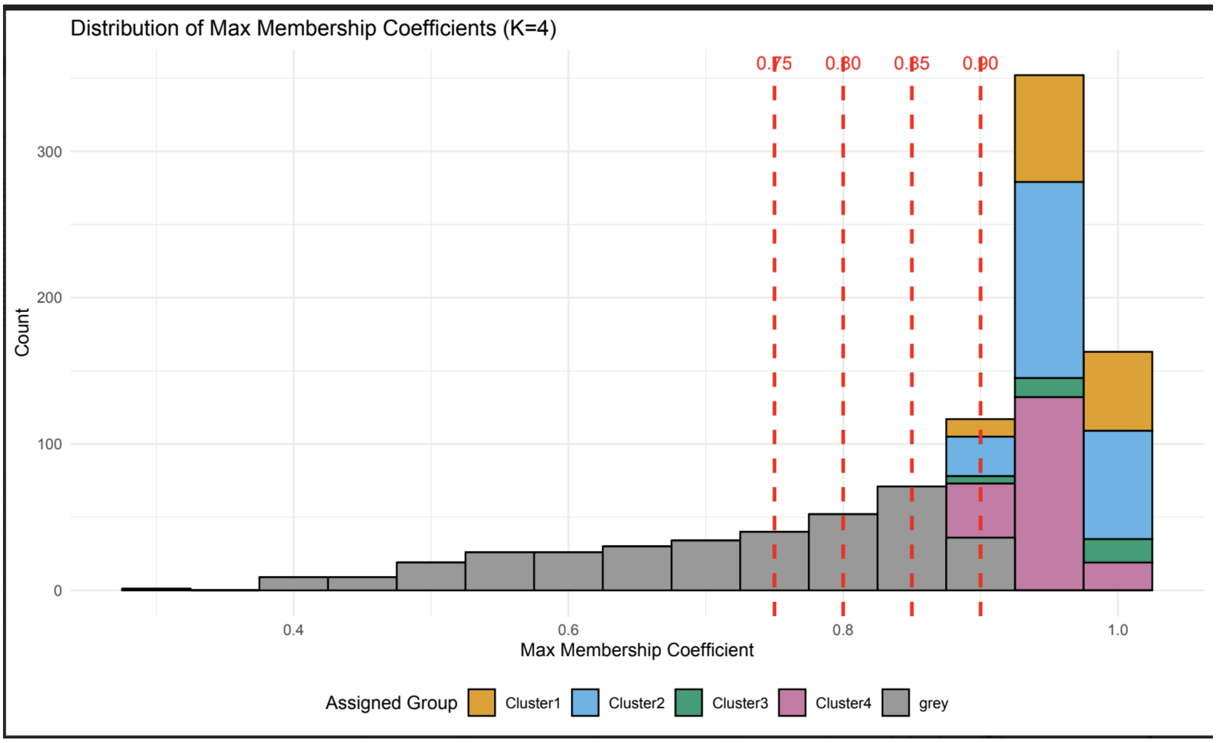


**Figure S3. Distribution of maximum membership coefficients for *K* = 4 in the full dataset of *Ficus* genotypes.**

This analysis consisted of *N*=949 *Ficus* genotypes with 884 *F. carica sensu stricto*, 38 *F. carica subsp. rupestris*, and 27 *F. colchica* individuals.

The histogram represents the distribution of individuals based on their highest STRUCTURE-inferred membership coefficient to one of four genetic clusters. Different colors indicate individuals assigned to specific clusters, while gray bars represent admixed individuals. Vertical red dashed lines indicate commonly used thresholds (e.g., 0.75, 0.80, 0.85, and 0.90) for distinguishing between "pure" individuals and admixed ones. The 0.90 threshold was used in this study to define "pure" cluster assignments.

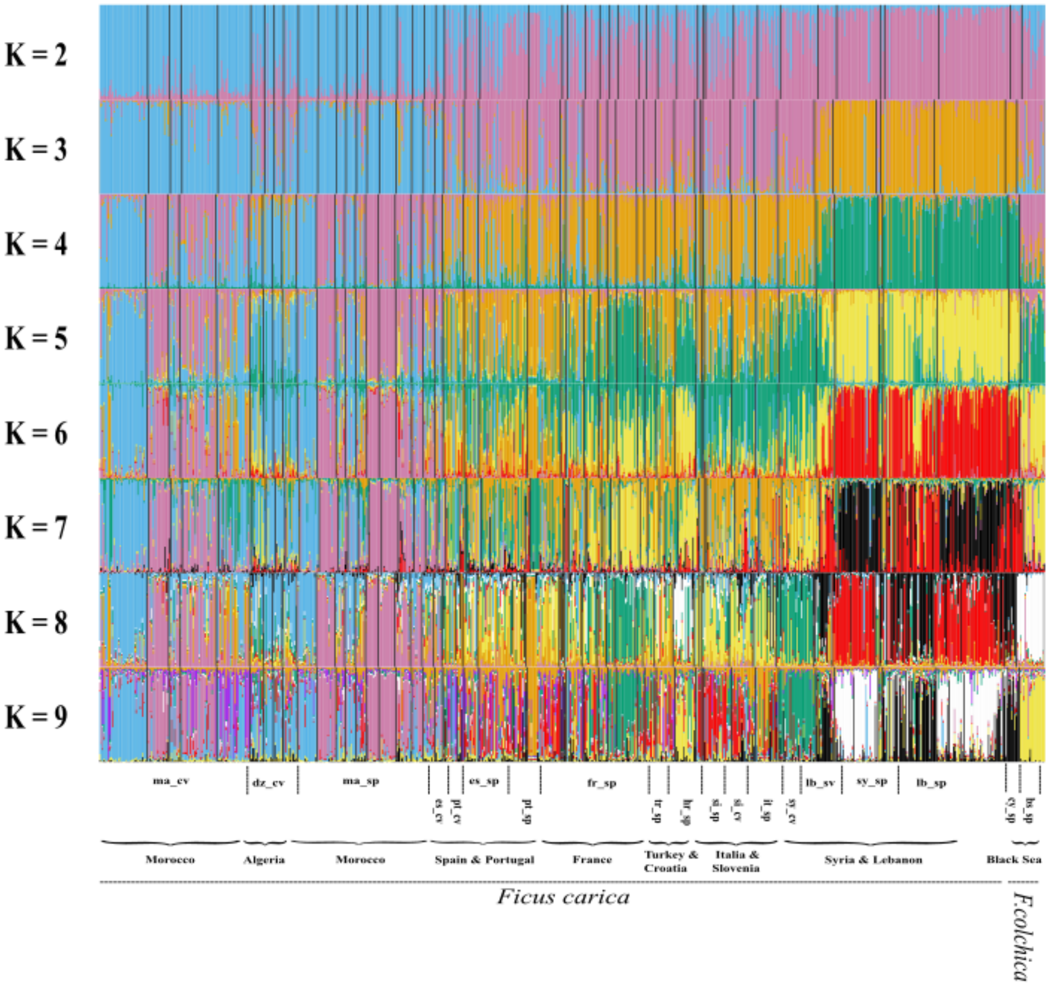


**Figure S4.** Proportions of ancestry for 911 *Ficus* genotypes assigned to K = 2 through K = 9 genetic clusters inferred by STRUCTURE.

This analysis was carried out on all *F. carica sensu stricto* and *F. colchica* samples, but excluding *F. rupestris* samples. Each individual is represented by a single vertical bar partitioned into K-colored segments, each segment corresponding to the proportion of the genome from an individual belonging to a given cluster. When multiple clustering solutions (modes) arose among replicate runs, the proportion of simulations supporting each mode is indicated.

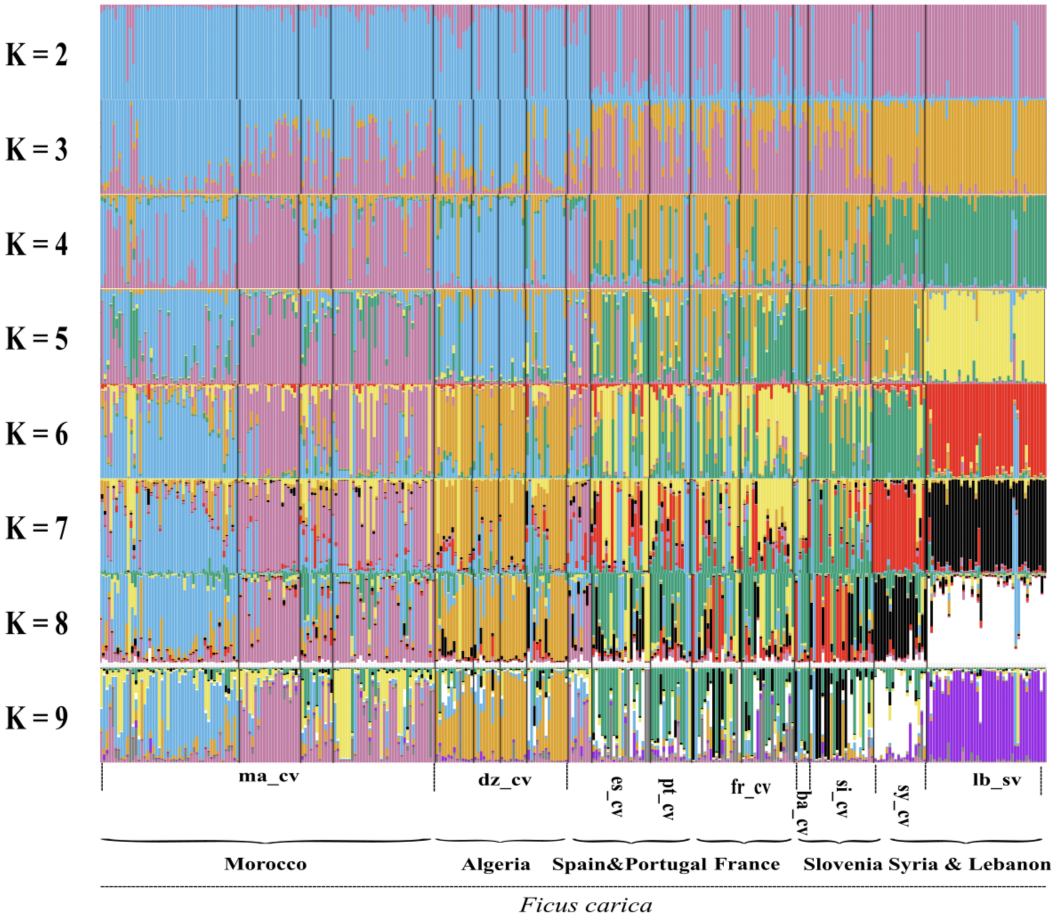


**Figure S5.** Proportions of ancestry for 911 *Ficus* genotypes assigned to K = 2 through K = 9 genetic clusters inferred by STRUCTURE.

This analysis was carried out on all *F. carica sensu stricto* (cultivated) samples, but excluding spontaneous *F. carica sensu stricto*, *F. carica subsp. rupestris*, and *F. colchica* samples. Each individual is represented by a single vertical bar, partitioned into K-colored segments, where each segment corresponds to the proportion of the genome from an individual belonging to a given cluster. When multiple clustering solutions (modes) arose among replicate runs, the proportion of simulations supporting each mode is indicated.

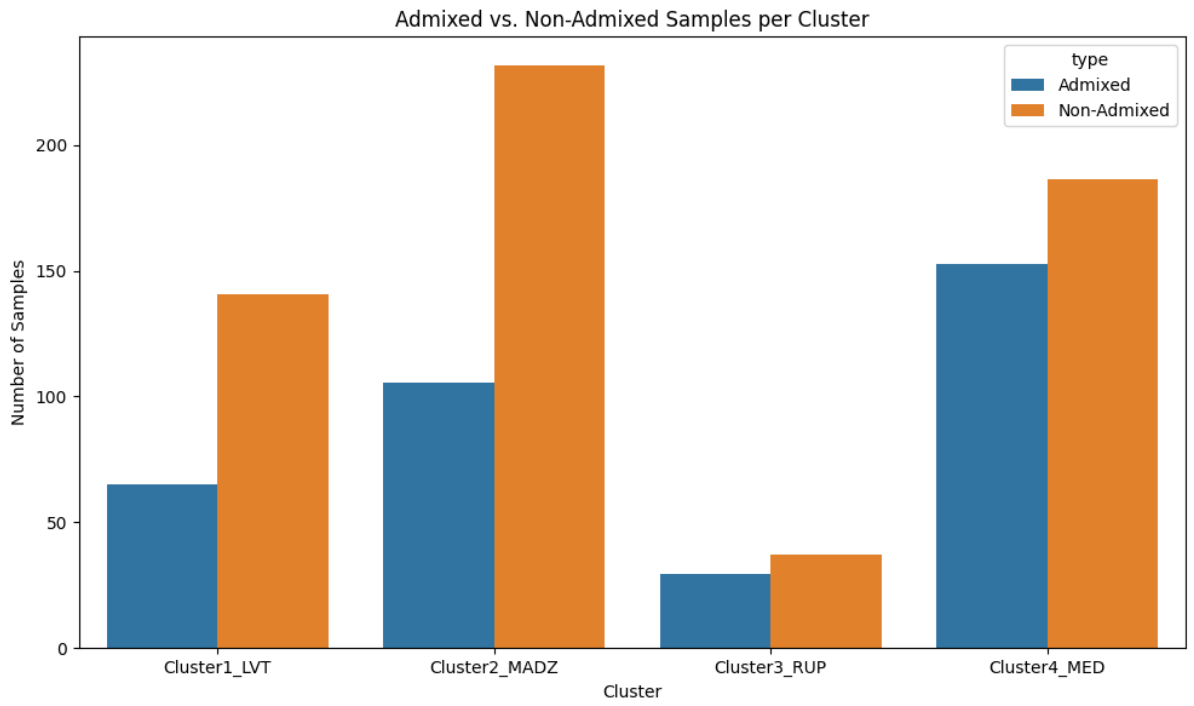


**Figure S6.** Number of admixed samples assigned to each of the four spontaneous or cultivated fig populations inferred at K=4.

Admixed individuals (i.e, individuals assigned <0.9 and > 0.55 to a given cluster) represented 37% of the dataset. The highest proportion of admixed individuals was observed in the cluster MED.

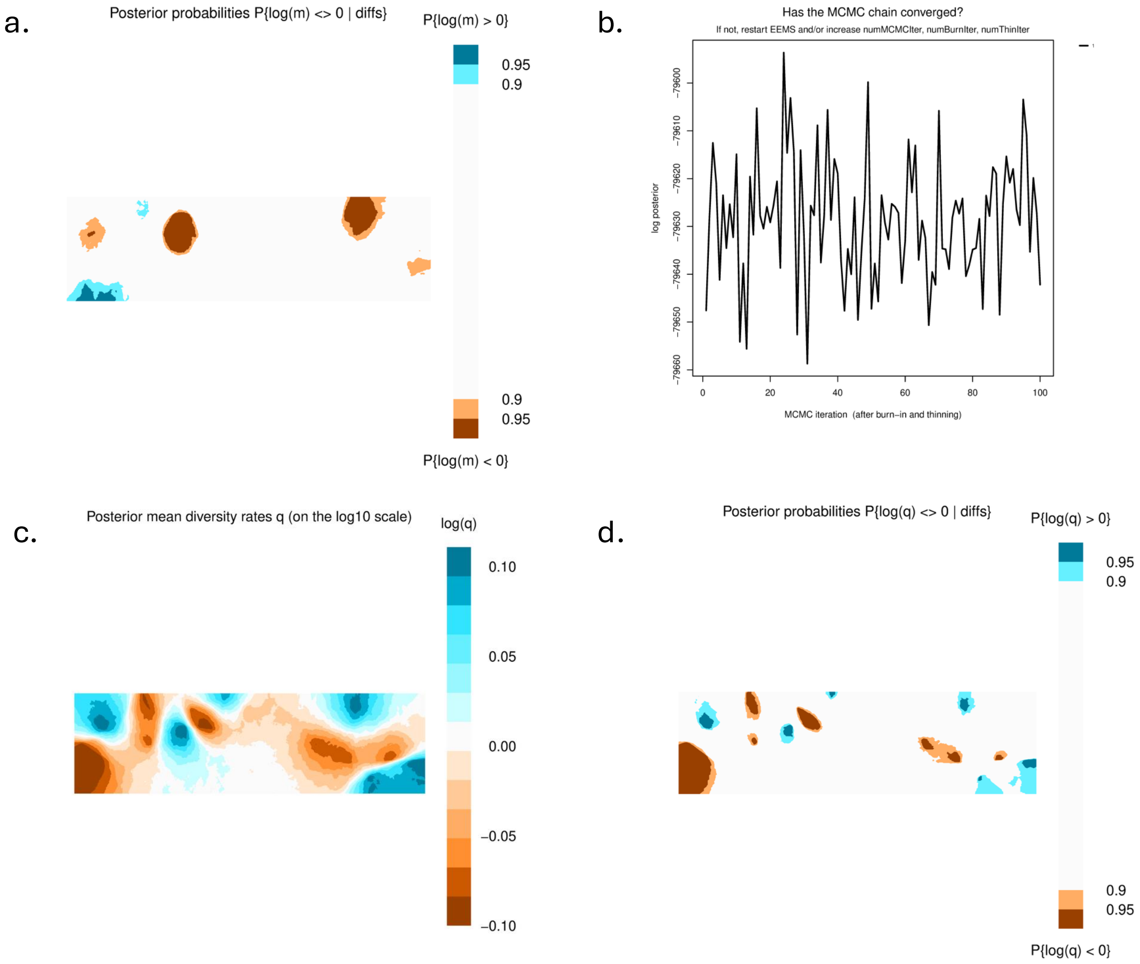


**Figure S7. Spatial patterns of migration and diversity inferred from EEMS (Estimated Effective Migration Surfaces).**

(a) Posterior probabilities for deviations in effective migration rates m, where brown and blue areas represent regions with significantly lower and higher migration rates, respectively [$P(|\log(m)| > 0)$]. (b) Markov chain Monte Carlo (MCMC) trace plot assessing convergence of the EEMS model across 100 iterations; stable fluctuations around the mean suggest proper chain convergence. (c) Posterior mean diversity rate q, $\log_{10}$-transformed, highlighting spatial variation in genetic diversity. (d) Posterior probabilities for deviations in the diversity rate q, with blue indicating higher than expected diversity and brown indicating lower diversity [$P(|\log(q)| > 0)$].

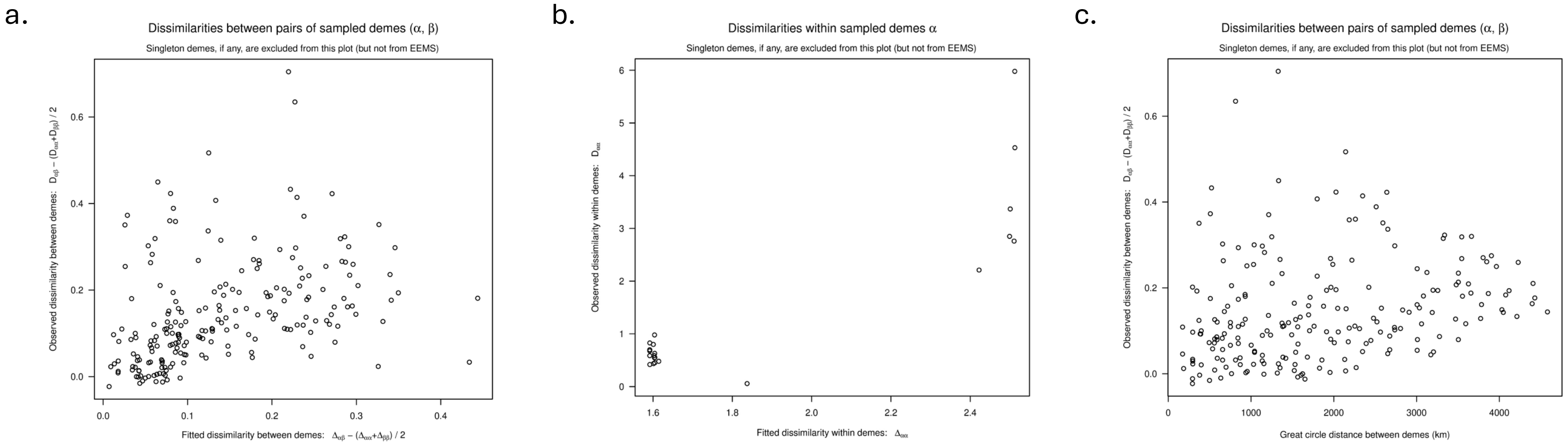


**Figure S8. Evaluation of the EEMS model fit through comparisons of observed and fitted genetic dissimilarities.**
(a) Scatterplot of observed versus fitted dissimilarities between pairs of sampled demes (D_{\alpha\beta}), showing moderate concordance between model predictions and empirical data. (b) Scatterplot of observed versus fitted dissimilarities within sampled demes (D_{\alpha\alpha}), highlighting overall fit at the deme level. (c) Scatterplot of observed genetic dissimilarities and great-circle geographical distances between demes (km), illustrating an isolation-by-distance (IBD) pattern. Although the EEMS model captures the broad trends in genetic structure, some variance remains unexplained, particularly across greater distances. These results support the presence of IBD and regional genetic structuring among fig populations, consistent with historical and ecological barriers to gene flow.
